# Effects of Hydrogen Peroxide on Organically Fertilized Hydroponic Lettuce (*Lactuca sativa* L.)

**DOI:** 10.1101/2021.01.01.425018

**Authors:** Vanessa Lau, Neil Mattson

**Affiliations:** School of Integrative Plant Sciences, Cornell University, Ithaca, NY 14853

**Keywords:** hydroponics, organic, biofilm, hydrogen peroxide, fertilizer

## Abstract

Hydroponic production typically uses conventional fertilizers and information is lacking on the use of organic hydroponic fertilizers. Development of biofilm is a common problem with organic hydroponics which can reduce dissolved oxygen availability to roots. One potential solution is the use of hydrogen peroxide, H_2_O_2_ which can reduce microbial populations and decomposes to form oxygen. However, information is lacking on the impact of hydrogen peroxide on hydroponic crops. The aim of this study was to determine the effects of H_2_O_2_ concentrations in deep water culture hydroponics by assessing how it affects plant size and yield in lettuce (Lactuca sativa L.) ‘Rouxai’. In this experiment, three different treatments consisting of a control without H_2_O_2_, and the application of 37.5 mg/L or 75 mg/L of hydrogen peroxide were added to aerated 4-L reservoirs that contained either organic (4-4-1) or inorganic nutrients (21-5-20), both applied at 150 mg·L^-1^ N. Three replicates for each treatment and each fertilizer were prepared resulting in a total of eighteen mini hydroponic containers each with one head of lettuce. When added to conventional fertilizers, concentrations of 37.5 mg/L and 75 mg/L of H_2_O_2_ led to stunted growth or death lettuce plants. However, when 37.5 mg/L of H_2_O_2_ was applied to organic fertilizers, the lettuce yield nearly matched that of the conventionally fertilized control, demonstrating that the application of H_2_O_2_ has the potential to make organic hydroponic fertilization a more viable method in the future.

## 1. Introduction

Marketed as a technologically revolutionary and sustainable way to grow produce, the employment of hydroponic methods in greenhouses and “plant factories” is gaining traction globally [1]. However, chemical fertilizers typically used in hydroponics are mined from nonrenewable, finite sources or rely on fossil fuels for production rendering them unsustainable [2,3]. Moreover, the disposal of chemically-fertilized waste water from these systems can leach into the environment and over time, degrade ecosystems as well as contaminate clean water sources [4,5].

An alternative to these conventional fertilizers is the use of organic fertilizers from plant and animal byproducts such as seaweed extract, manure, or hydrolyzed fish emulsion [6] which require the development of microbial communities to mineralize complex organic compounds to make them plant available [7,8]. Drawbacks of organic fertilizers include variable, significantly reduced yield [9–11] which may be attributed to: unstable microbial activity, difficulty supplying the proper proportion of nutrients, high pH as well as the development of biofilm in the organically fertilized hydroponic reservoirs [12]. Regarding biofilm, it is believed that the suspended organic matter, which can develop on plant roots as well as clog pumps/recirculation lines, reduce oxygen and nutrient uptake by roots as well as deplete oxygen levels [13–15].

There is some anecdotal evidence that the addition of hydrogen peroxide (H_2_O_2_) to organically fertilized reservoirs may help reduce the development of biofilm and improve performance of organic hydroponics [16,17]. H_2_O_2_ is an unstable oxidizing agent most commonly used as an inexpensive, household disinfectant and bleaching agent [18]. Byproducts produced by the decomposition of H_2_O_2_ are H_2_O and O_2_. The released O_2_ can increase the dissolved oxygen concentration in the root zone and may also help reduce oxygen losses to biofilm and microbial respiration. Though the application of H_2_O_2_ is thought to help increase DO concentrations within the reservoir, in conventional hydroponics lettuce studies did not previously show a positive benefit of DO at or above 25% of saturation (2.1 mg·L^-1^) on shoot or root biomass [19]. Possible benefits of H_2_O_2_ in an organic hydroponic system could be to increase dissolved oxygen concentration if they fall below this low threshold or alternatively may be due to H_2_O_2_’s disinfectant properties.

However, currently there is no work in the scientific literature on H_2_O_2_ use in conventional or organic hydroponic systems, including no commonly recognized concentration or range of H_2_O_2_ to add to hydroponic systems. Excess H_2_O_2_ can also harm plant root systems in hydroponics [16,20,21] and information is lacking on the concentration that damages hydroponic crops, including lettuce. Currently suggested H_2_O_2_ practices vary greatly among hobbyists and are typically determined on a trial and error basis. With little to no scientifically backed information available on the topic, this study aims to explore the usage of H_2_O_2_ on dissolved oxygen in the root zone and its effects on yield in conventionally and organically fertilized lettuce heads.

## 2. Materials and Methods

### 2.1. Experimental Seedling Preparation

The experiments were conducted at the Cornell University Kenneth Post Laboratory greenhouses in Ithaca, New York at room temperature (22°C) with ambient lighting, with the first crop cycle taking place from November through December 2018 and the second crop cycle taking place March to April 2019. A flat with 1.5 x 1.5 inch rockwool plugs was soaked in reverse osmosis water for approximately 10 minutes and then was drained and placed on a plastic flat. Each rockwool cell was seeded with one pelleted (*Lactuca sativa* L.) ‘Rouxai’ seed (Johnny’s Seeds, Fairfield, ME) and was allowed to germinate in a seedling production area in the greenhouse under 18-hour lighting from high pressure sodium (HPS) lamps. The seedlings were watered daily with fertilized water (Jack’s Professional LX 21-5-20 All Purpose Water-Soluble Fertilizer supplemented with magnesium sulfate so as to supply 30 ppm Mg). Seedlings were transplanted into 4-L hydroponic containers after 21 days.

### 2.2. Treatment Setup

After 21 days when the seedlings had 3 to 4 true leaves, eighteen individual 4-L buckets were prepared for the experiment. Each bucket was filled near to capacity with reverse osmosis water. The conventional fertilizer, Jack’s Professional LX 21-5-20 All Purpose Water-Soluble Fertilizer (JR Peters Inc., Allentown, PA) with magnesium sulfate was applied to half of the buckets, the other half of the buckets were fertilized with the organic fertilizer, Drammatic One 4-4-1 Fish Emulsion (Dramm Corporation, Manitowoc, WI) and in both cases the fertilizers were added to supply an electrical conductivity (EC) of 1.5-1.7 dS/m. H_2_O_2_ was added to the buckets according to the treatments in Table 1 to create 3 replications of each treatment.

**Table 1.** Names and descriptions of experimental treatments

| Treatment | Description |
| --- | --- |
| CONV_0 | Conventionally fertilized control |
| CONV_37.5 | Conventionally fertilized with 37.5 mg/L H<sub>2</sub>O<sub>2</sub> |
| CONV_75 | Conventionally fertilized with 75 mg/L H<sub>2</sub>O<sub>2</sub> |
| ORG_0 | Organically fertilized control |
| ORG_37.5 | Organically fertilized with 37.5 mg/L H<sub>2</sub>O<sub>2</sub> |
| ORG_75 | Organically fertilized with 75 mg/L H<sub>2</sub>O<sub>2</sub> |

The buckets were arranged on a bench in the greenhouse in 3 rows of 6 buckets spaced approximately 30 cm apart and each bucket was individually aerated with an airstone placed near the rootzone that was powered by air pumps (GH2716, General Hydroponics, Santa Rosa, CA) as shown in Figure 1 and were randomly arranged by treatment.

**Figure 1.**
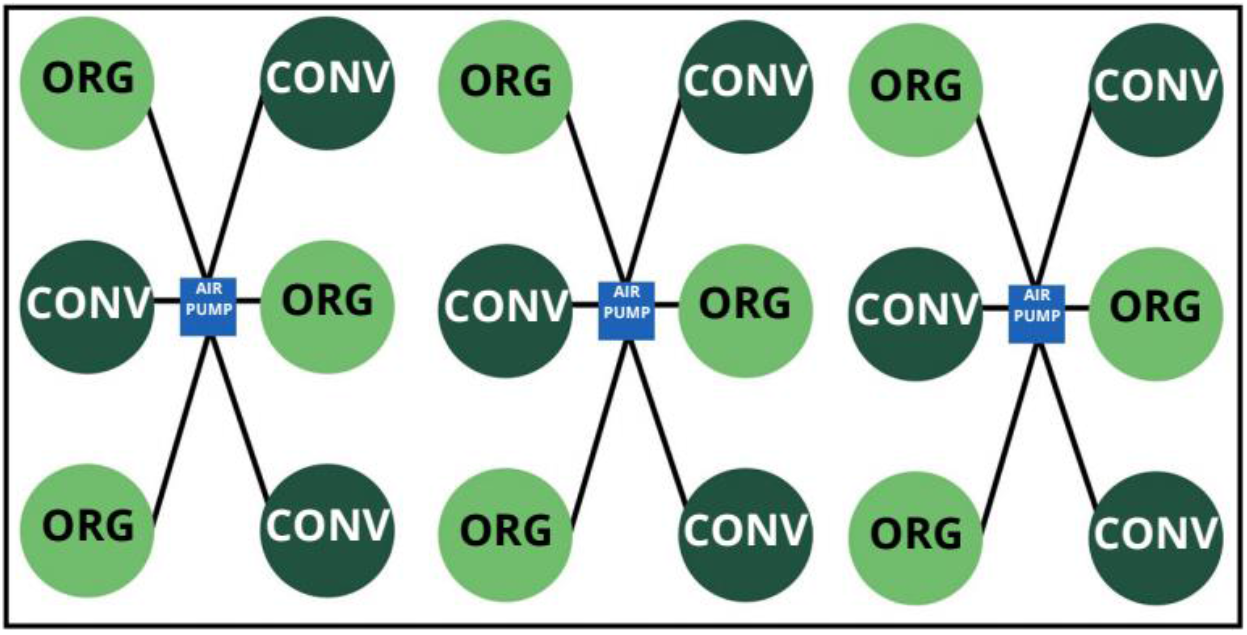
Illustration of experimental set up

The electrical conductivity (EC) of each bucket was measured using an EcoTestr CTS meter (Oakton, Vernon Hills, IL) and was adjusted to 1.5-1.7 dS/m. Additionally, the pH levels were measured using an EcoTestr pH2+ meter (Oakton, Vernon Hills, IL) and were adjusted to 5.5-6 with either nitric acid (1M HNO_3_) to lower the pH or potassium hydroxide (1M KOH) to bring pH levels up.

18 uniform lettuce seedlings were selected from the seedling production area and each plant was placed in the 1” diameter hole drilled into the center of each lid so that the rockwool plugs fit snugly. The buckets were checked to make sure the water levels were high enough to reach the plants’ roots. Dissolved oxygen (DO) measurements were taken using a YSI Pro20 Dissolved Oxygen Meter (Xylem Inc., Yellow Springs, OH) and results are expressed on a percent of DO saturation at the recorded water temperature.

### 2.3. Experimental Protocol & Data Collection

Each day, EC and pH were adjusted and maintained between the target values (1.5-1.7 dS/m and pH 5.5-6) using the respective tools and reservoir water levels were topped-off daily to maintain 4-L of nutrient solution. DO measurements were recorded daily.

Every 3 days, hydrogen peroxide was replenished according to the treatment prior to the DO measurements for that day. This was done for 17 days and on the 18th, lettuce was harvested by severing the plant where the plant stalk met the rockwool plugs and final fresh weight was recorded.

## 3. Results

### 3.1. Dissolved Oxygen

In the Fall trial, dissolved oxygen (DO) levels were recorded each day to track the degradation of H_2_O_2_ within the rootzone. DO was added to the hydroponic containers 3 times weekly (represented by days 0, 3, 6, 12, 15…) on Figure 2. Steep increases in DO levels typically represent days in which H_2_O_2_ was added to the reservoirs. It was noted that on average, organically fertilized treatments saw more drastic swings in DO levels than their conventional counterparts and that over time, the application of H_2_O_2_ had lessened effects on DO levels. Conventional fertilizer with 75 mg/L H_2_O_2_ led to the greatest sustained levels of DO, in the organic treatments DO level degraded more quickly after each addition.

**Figure 2.**
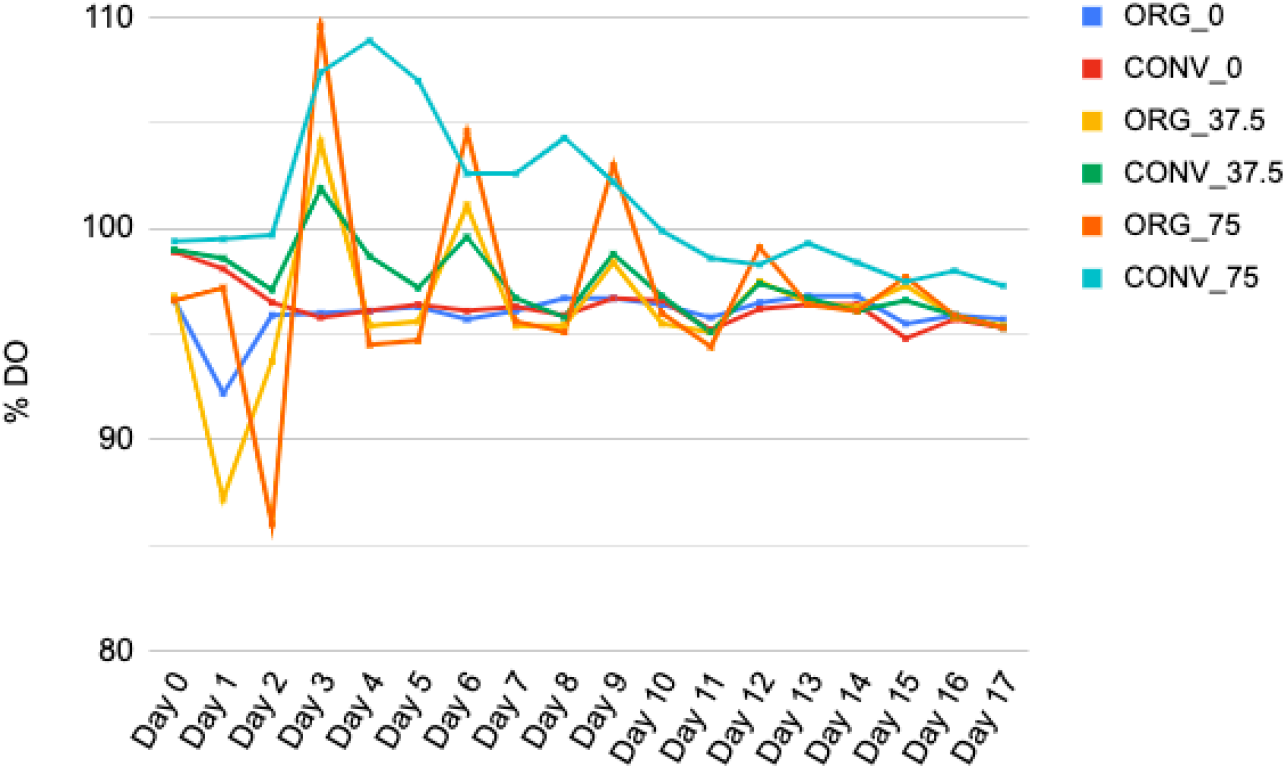
Percent dissolved oxygen levels taken within lettuce plant root zones in the fertilizer and H_2_O_2_ treatments in the Fall trial. Data are means of 3 plants per treatment combination per day.

### 3.2. Fresh Weight

The plants in the CONV_0 treatment had the highest mean value, with no statistically significant difference between this treatment and organic or conventional treatments with 37.5 mg/L H_2_O_2_ (Figure 3). The conventionally fertilized control yield was nearly double that of the organically fertilized with 0 mg/L H_2_O_2_. With the lower application of H_2_O_2_, however, ORG_37.5 had a statistically similar FW as CONV_37.5 and CONV_0. At the greater application of 75 mg/L H_2_O_2_, both ORG_75 and CONV_75 yield decreased with the CONV_75 plants dying and therefore having a lower FW than all other treatments. Across treatments, this shows that application of H_2_O_2_ had a negative impact on CONV but low doses (37.5 mg/L) actually increased yield of organic treatments to the point that it was not significantly different from conventional fertilizer.

**Figure 3.**
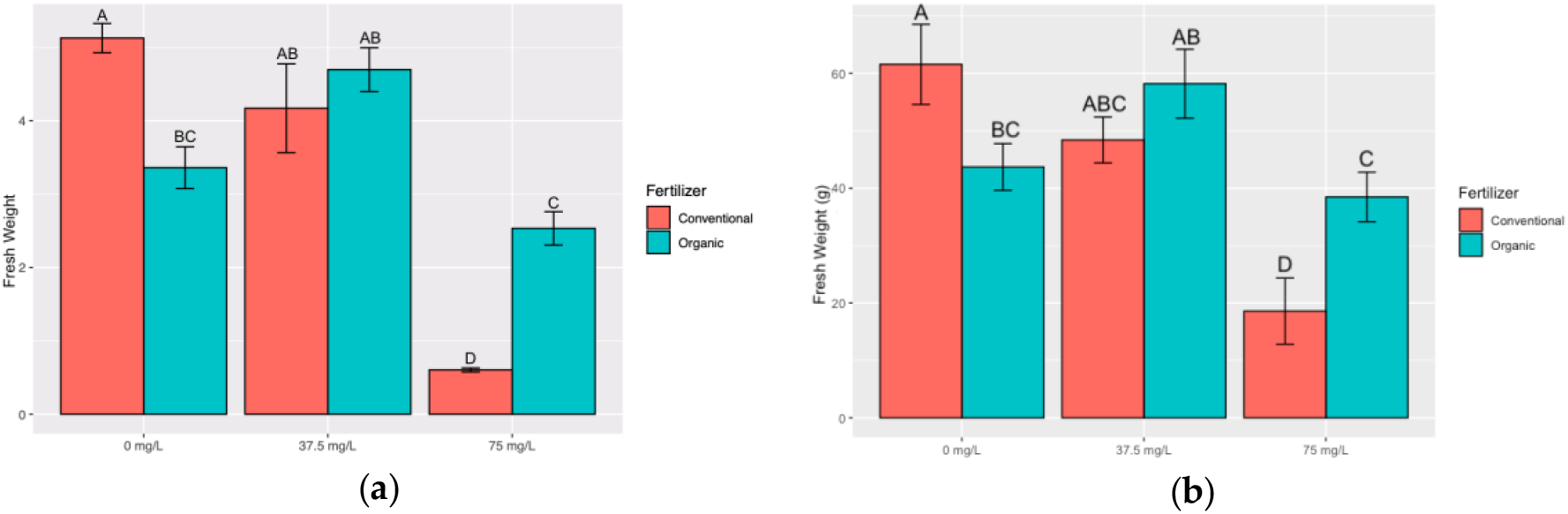
Fresh weight in grams of lettuce plants in response to organic and conventional fertilizer and addition of H_2_O_2_ in the Fall (**a**) and Spring (**b**) trial. Data are means ± SE of 3 plants per treatment combination. Letters represent mean separation comparison using Tukey’s HSD (alpha=0.05).

During the Spring 2019 trial, trends between treatments remained similar confirming the results from the Fall 2018 trial (Figure 3). However, overall fresh weight was greater in the Spring trial due to greater ambient light. Additionally, though their growth was severely stunted, the plants grown with the CONV_75 and ORG_75 treatments with high doses of H_2_O_2_ did not die.

### 3.3. Root Length

In the Fall trial plants in the CONV_0 and ORG_0 treatment had the highest mean root length (Figure 4). The application of H_2_O_2_ dramatically decreased root length for both fertilizer treatments but the effects were more dramatic for conventionally fertilized treatments.

**Figure 4.**
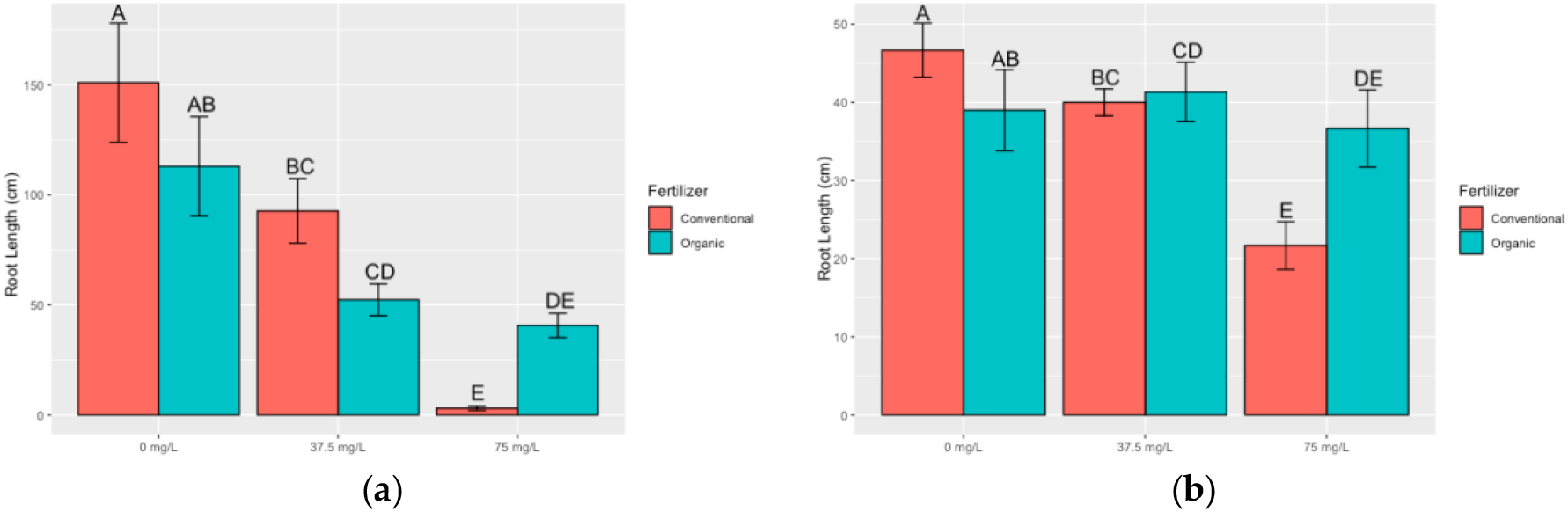
Root length of lettuce plants in response to organic and conventional fertilizer and addition of hydrogen peroxide in the Fall (**a**) and Spring (**b**) trial. Data are means ± SE of 3 plants per treatment combination. Letters represent mean separation comparison using Tukey’s HSD (alpha=0.05).

In the Spring trial similar patterns were found. There was no difference in root length between the conventional and organic controls, but as H_2_O_2_ treatments increased, root length dramatically decreased, especially in the conventionally fertilized treatments.

### 3.4. Leaf Width and Plant Height

For the Fall trial, data was collected on leaf width and plant height. Due to time constraints these parameters were not collected in the Spring trial. For leaf width, the only statistically significant difference was that CONV_75 had a smaller leaf width (about half the size) as all other treatments (Figure 5).

**Figure 5.**
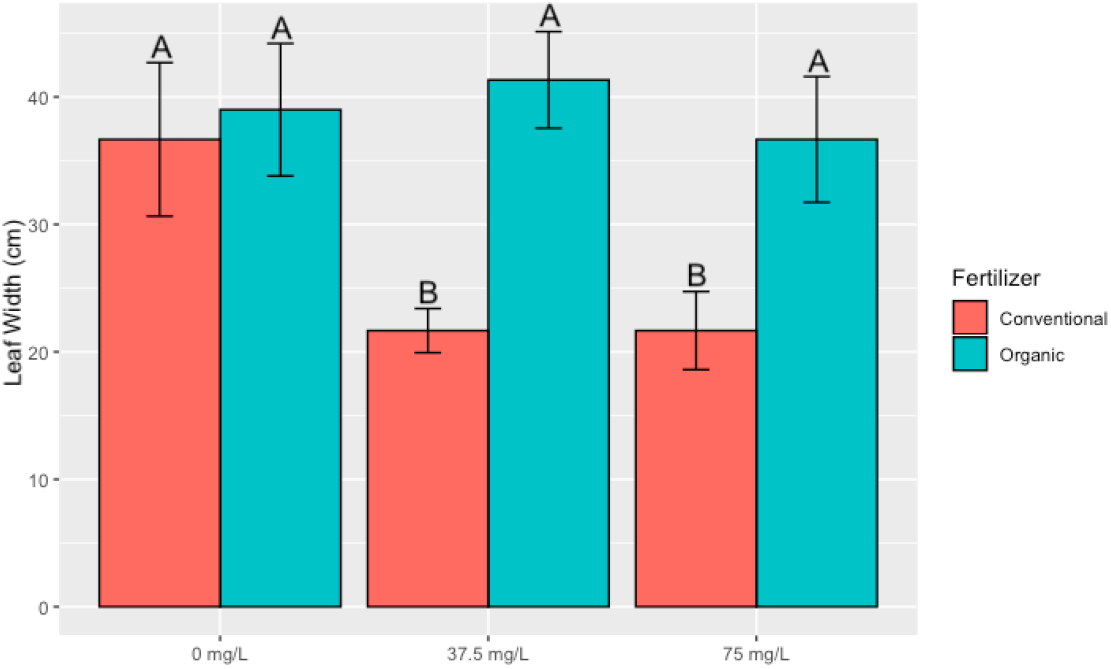
Leaf width of lettuce plants in response to organic and conventional fertilizer and addition of hydrogen peroxide in the Fall trial. Data are means ± SE of 3 plants per treatment combination. Letters represent mean separation comparison using Tukey’s HSD (alpha=0.05).

Likewise, for plant height, similar trends were found whereby height of conventionally fertilized plants at 75 mg/L H_2_O_2_ was dramatically smaller than other treatments. For organic fertilization, the plants at 75 mg/L H_2_O_2_ were shorter than conventional plants at 37.5 mg/L H_2_O_2_ (Figure 5).

**Figure 6.**
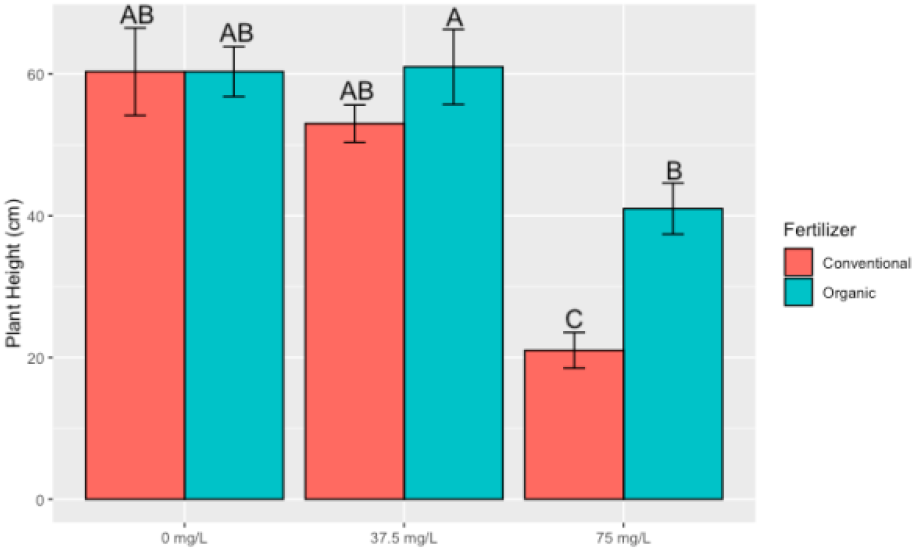
Plant height of lettuce plants in response to organic and conventional fertilizer and addition of hydrogen peroxide in the Fall trial. Data are means ± SE of 3 plants per treatment combination. Letters represent mean separation comparison using Tukey’s HSD (alpha=0.05).

## 4. Discussion

The results of this study show the potential for the integration of H_2_O_2_ with organic fertilizers to optimize hydroponic lettuce yield. Without hydrogen peroxide, our study found that organic fertilization performed poorer (fresh weight, root length) than conventional fertilizer. Our findings with organic fertilization performance (without H_2_O_2_) are similar to those reported by Atkin & Nichols [11]. For example, Atkin & Nichols also tested hydrolyzed fish emulsion based fertilizer and found that organic hydroponic lettuce had approximately 55% lower fresh weight than conventional. Results from the conventional and organic controls matched the % yield ranges of other studies [11,22,23]

H_2_O_2_ additions reduced the performance of conventionally fertilized plants. H_2_O_2_ works by decomposing into an unstable free radical oxygen molecule which can destroy biotic cell tissue. As such, H_2_O_2_ has the potential to indiscriminately damage healthy living root tissue, consequently reducing fresh weight of lettuce heads in higher doses. Root damage may be the result of this phytotoxicity [24,25,26]. However, when H_2_O_2_ at 37.5 mg/L was added to organic fertilizer the organic fertilizer performed as well as conventional fertilizer without H_2_O_2_. Whereas H_2_O_2_ at 37.5 mg/L or 75 mg/L. We hypothesize that the lack of negative impacts of 1.25 mL/L H_2_O_2_ in the organic fertilizer treatment were due to the effect of biofilm present in the rootzone - the free radical oxygen molecules may have disrupting biofilm matter thereby leading to less damage to roots. Since the conventionally fertilized reservoirs did not contain visible biofilm, this may have led to higher levels of root damage, and subsequently, lower lettuce fresh weight.

Moreso, while the effects of the application of H_2_O_2_ had a visible impact on the plant material, there were no visible reductions in biofilm development among the organic treatments. This indicates that while it may be effective in increasing yield, manual disinfection of hydroponic systems would still be needed in between growth cycles to clear out the biofilm that may clog and stick to surfaces. Though future research would be needed to quantify the impact of H_2_O_2_ on biofilm in hydroponics.

## 5. Conclusions

Overall it was found that the use of 37.5 mg/L H_2_O_2_ could lead to effective performance of plants receiving organic hydroponics. More research is needed to determine if the response is due to 1) the increase in dissolved oxygen content of the root-zone as H_2_O_2_ disassociated, 2) the effects of H_2_O_2_ on biofilm development (i.e. injuring biofilm but allowing for greater nutrient or dissolved oxygen access by roots), or 3) the effect of H_2_O_2_ on the chemical makeup (nutrient availability) or the organic hydroponic nutrient solution. Future research should seek to understand the optimum concentration and reapplication rate of H_2_O_2_ in organic hydroponic fertilization and well as to understand the mechanism for the response so that we have a greater understanding of effective organic hydroponic fertilization strategies.

## Author Contributions

Conceptualization, V.L.; methodology, V.L. and N.M.; formal analysis, V.L. and N.M..; writing—original draft preparation, V.L.; writing—review and editing, N.M.; supervision, N.M.; project administration, N.M. All authors have read and agreed to the published version of the manuscript.

## Acknowledgments

I wish to express my sincere gratitude to Matthew Moghaddam and Nicholas Kaczmar for their valuable technical support. Without their assistance, this paper and the research behind it would not have been possible.

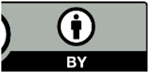 20 by the authors. Submitted for possible open access publication under the terms and conditions of the Creative Commons Attribution (CC BY) license (http://creativecommons.org/licenses/by/4.0/).

